# Incremental BLAST: incremental addition of new sequence databases through e-value correction

**DOI:** 10.1101/476218

**Authors:** Sajal Dash, Sarthok Rahman, Heather M. Hines, Wu-chun Feng

**Affiliations:** Department of Computer Science, Virginia Tech, Blacksburg, VA, USA; Department of Biology, The Pennsylvania State University, University Park, PA, USA; Department of Entomology, The Pennsylvania State University, University Park, PA, USA; Department of Electrical and Computer Engineering, Virginia Tech, Blacksburg, VA, USA

**Keywords:** Sequence similarity search, BLAST, e-value correction, taxon-specific BLAST, incremental search, local alignment

## Abstract

**Motivation:** Search results from local alignment search tools use statistical parameters sensitive to the size of the database. NCBI BLAST, for example, reports important matches using similarity scores and expect or e-values calculated against database size. Over the course of an investigation, the database grows and the best matches may change. To update the results of a sequence similarity search to find the most optimal hits, bioinformaticians must rerun the BLAST search against the entire database; this translates into irredeemable spent time, money, and computational resources.

**Results:** We develop an efficient way to redeem spent BLAST search effort by introducing the Incremental BLAST. This tool makes use of the previous BLAST search results as it conducts new searches on only the incremental part of the database, recomputes statistical metrics such as e-values and combines these two sets of results to produce updated results. We develop statistics for correcting e-values of any BLAST result against any arbitrary sequence database. The experimental results and accuracy analysis demonstrate that Incremental BLAST can provide search results identical to NCBI BLAST at a significantly reduced computational cost. We apply three case studies to showcase different use cases where Incremental BLAST can make biological discovery more efficiently at a reduced cost. This tool can be used to update sequence blasts during the course of genomic and transcriptomic projects, such as in re-annotation projects, and to conduct incremental addition of taxon-specific sequences to a BLAST database. Incremental BLAST performs (1 + *δ*)*/δ* times faster than NCBI BLAST for *δ* fraction of database growth.

**Availability:** Incremental BLAST is available at https://bitbucket.org/sajal000/incremental-blast.

**Supplementary information:** Supplementary data are available at https://bitbucket.org/sajal000/incremental-blast

## 1 Introduction

Utilization of a sequence similarity search tool is a central step in most bioinformatics research investigating biological or structural functions of nucleotide or protein sequences. *BLAST*, (Basic Local Alignment Search Tool) (Altschul et al. 1990) is a widely used (74,000+ citations, October 2018) sequence alignment tool capable of conducting a sequence similarity search for a sequence of interest against a curated sequence database. BLAST relies on a heuristic approach for searching and provides results based on the identification of regions through seed-and-extend based local alignment which could either be implemented using a web interface (Johnson et al. 2008) or a set of standalone command line tools maintained by NCBI (Camacho et al. 2009). It uses a statistical threshold, an expect value or e-value, to infer homologous sequences from a curated database. BLAST is extensively used for identification of unknown sequences, detection of candidate genes, and during the annotation of assembled genomes and transcriptomes.

Sequencing data stored in the NCBI database has expanded astronomically over the years, reportedly doubling in the number of bases submitted to Genbank every year over the last three decades (1982-present; e.g., Figure 1). Cheaper sequencing technology (Figure 1), the democratization of HPC through commercial cloud platforms, improvement of genomic/transcriptomic assemblers and pipelines, and efforts to sequence many new taxa (e.g., Earth BioGenome Project (Lewin et al. 2018), The i5k Initiative (i5K Consortium 2013), BAT 1K (Skibba 2016), The Genome 10K Project (Koepfli et al. 2015)) are some of the major reasons facilitating this growth. Fast-accumulating sequences in NCBI curated databases has a profound impact on the computational efforts required to perform sequence similarity search.

**Figure 1:**
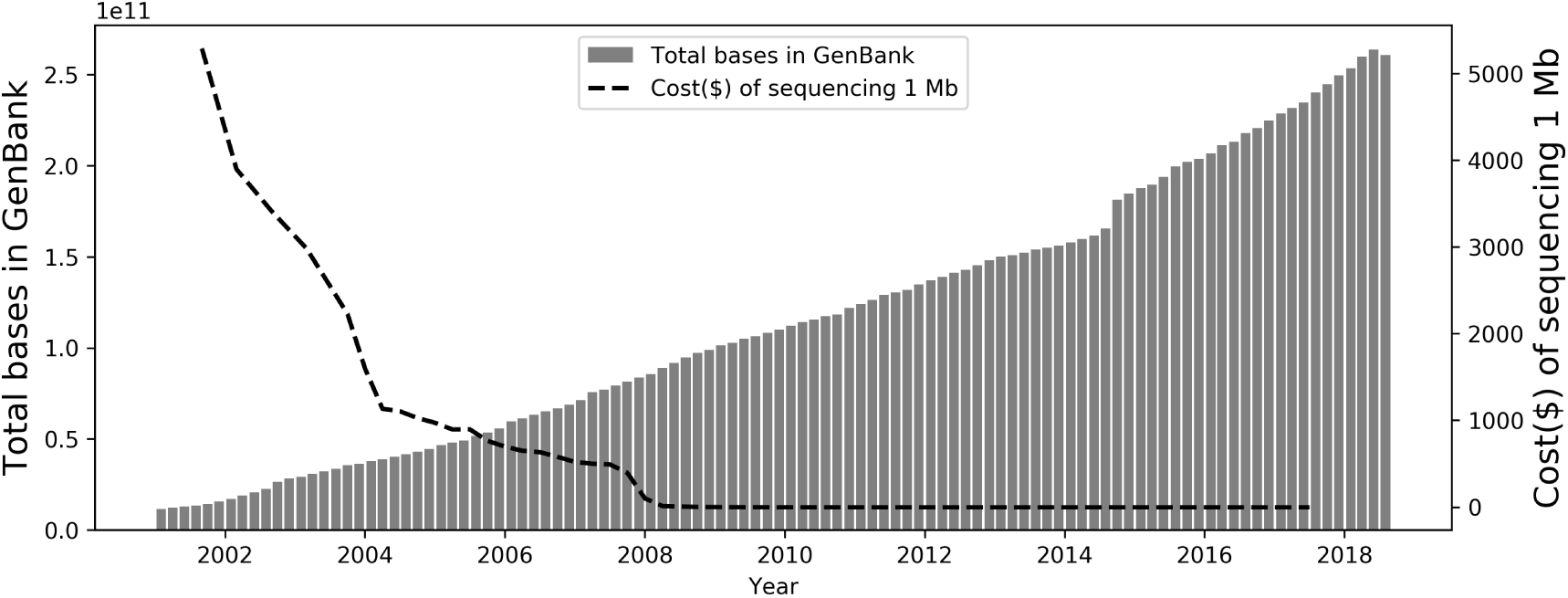
Increasing GenBank database size (Ncbi.nlm.nih.gov 2018 (accessed September 15, 2018) follows a decreasing trend in sequencing cost (Wetterstrand 2018 (accessed September 15, 2018).

It has been an active area of bioinformatics research to provide fast and biologically valuable sequence alignment tools to deal with this ever-growing database. Some sequence alignment programs have tried to make some algorithmic improvements (HMMER (Eddy 1998), DIAMOND (Buchfink et al. 2014)) while others have focused on improving parallelization and taking advantages of new HPC platforms and programming paradigms, including cuBLASTP (Zhang et al. 2014), muBLASTP (Zhang et al. 2016), mpiBLAST (Darling et al. 2003), SparkBLAST (de Castro et al. 2017) BLAST tools. All of these tools provide similar output to NCBI BLAST at an improved computational speed. Other BLAST tools provide convenience factors of BLAST usage such as NOBLAST (Lagnel et al. 2009), which offers new options to limit the number of sequences to search, and JAMBLAST (Lagnel et al. 2009) which provides visualization tools for NOBLAST output.

BLAST is computationally expensive to run, with computational time impacted by the number of queries and reference database size. Genome sequencing and annotation projects can be fairly long-term projects that require updates mid-project and regular annotation updates. However, for such updates, sequence similarity search steps have to be executed from scratch as search results from BLAST use metrics similarity scores and e-values that depend on the size of the database, which is continually changing. For this reason, it is required to rerun the entire search which renders previous searches useless, translating to irredeemable time, money, and computational resources.

For bioinformatics projects requiring large-scale sequence alignment task, such as those involving many transcriptomes from many taxa, the computational burden can be especially prohibitive, a problem that could be solved through performing iterative taxon-specific searches rather than conducting BLAST to the entire non-redundant database. However, such an approach has been historically difficult as one would need to standardize e-values when iteratively adding new databases to find the optimal identity of each query.

Here we introduce Incremental BLAST which provides an efficient solution to e-value correction and allows the merging of results from two separate databases, thus allowing recycling of previous results, which subsequently saves both time and money. It also enables taxon-specific BLAST search, including incremental addition of specific biologically-relevant taxa to BLAST databases with subsequent merging. Our approach will improve time-savings for large-scale projects, and can make it easier to sort hits by taxon. The Incremental BLAST tool is easy-to-use, simply involving the NCBI BLAST original program and few python modules, and it works with different versions of NCBI BLAST command line tools.

## 2 Background and Related Work

### 2.1 Core concepts of BLAST Result: Hit, HSP, Score, E-value

When a BLAST search is performed against a sequence database with a query sequence, the best matching sequences from the target databases are returned by the BLAST program. These best matching sequences are *hit*s. Between the query and a hit sequence, there are many pairwise local matches. They are called high scoring pairs or *HSP*. One hit consists of many HSPs. HSPs are scored using some statistical matrices by comparing aligned symbols. The score for a hit is the score of highest scoring HSP that belongs to that hit. The e-value for an HSP is computed using the score, the database size, and other statistical parameters. The reported e-value of a hit is the e-value of the HSP with the lowest e-value.

### 2.2 BLAST Statistics for E-value computation

BLAST programs use two different kinds of statistics for e-value computation: Karlin-Altschul statistics and Spouge statistics.

#### 2.2.1 Karlin-Altschul Statistics

Gumbel EVD is often used to approximate the distributions of the maxima of long sequences of random variables, in this case, the distributions of the HSPs. The Gumbel EVD states that the probability of a score *x* greater than or equal to *S* is *p*(*x ≥ S*) = 1 *− e*^*−λ*(*S−µ*)^. Here *λ* is the scale parameter, and *µ* is the location parameter. Karlin and Altschul established a statistical theory about local alignment statistics using Gumbel EVD under certain assumptions to derive the formula for E-value *E*,

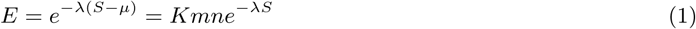

which is the famous Karlin-Altschul equation (Altschul et al. 1990).

**Edge Effect** Karlin-Altschul derives e-value statistics under the assumption that the sequence-lengths are infinite. This does not hold with the introduction of the new length parameters *m* and *n*. The parameters *m* and *n* here are called the effective length of the query and the database. They are introduced to compensate for the edge effect of alignments, which occurs at the end of the query sequence or the database sequence, where there may not be enough sequence space to construct an optimal alignment. So, the effective lengths are computed using a length adjustment *l*. Here, *m* = *m_a_ − l, n* = *n_a_ − N* × *l, l* = ln (*K × m × n*)*/H*. Here *N* is the number of sequences in the database, and *H* is the entropy of the scoring system. The length adjustment *l* satisfies equation 2.

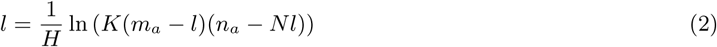

#### 2.2.2 Spouge Statistics

Spouge statistics (Park et al. 2012) is developed on the Karlin-Altschul formula. Instead of computing length adjustment *l* and then using it to compute the effective length of the database and query a finite size correction (FSC) is applied.

Finite size correction (Park et al. 2012) was introduced since version 2.2.26. Instead of estimating *l*, FSC estimates

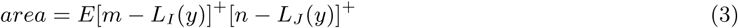

as a measure of (*m − l*)(*n − Nl*). Here, *I, J* are two sequences to be compared. *L_I_* (*y*) is the distribution of the length required to attain a score of *y* or more. Equation 3 is practically computed by approximating the distribution of *〈L_I_*(*y*)*, L_J_* (*y*)*〉*.

There is a range of statistical parameters which are used to compute the area. These parameters don’t depend on the length of the database or the query. However, in actual BLAST implementations using Spouge statistics, the formula is modified to include a database scale factor. The database scale factor is calculated using the formula,

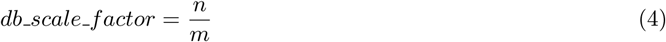

For a given HSP with score *S*, e-value is calculated using

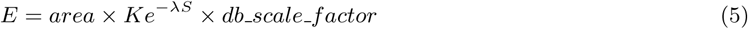

#### 2.2.3 Different Statistics used by BLAST Programs

Table 1 shows various statistics used by various BLAST programs.

**Table 1:** Both Spouge and Karlin-Altschul statistics are used by various NCBI BLAST programs.

| BLAST Program | Function | E-value Statistics |
| --- | --- | --- |
| blastn | Search Nucleotide sequences with a Nucleotide query | Karlin-Altschul |
| blastp | Search protein database with a protein query | Spouge |
| blastx | Search protein database with a Nucleotide query translated to protein | Spouge |
| tblastx | Search a translated Nucleotide database with a translated Nucleotide query | Karlin-Altschul |
| tblastn | Search a translated Nucleotide database with a protein sequence | Spouge |

### 2.3 Existing e-value correction software and their features

#### 2.3.1 mpiBLAST

mpiBLAST (Darling et al. 2003) is a parallel implementation of NCBI BLAST on the cluster. It segments the database, ports the segments into different nodes of a cluster, and runs parallel BLAST search jobs against database segments on different nodes. Once the parallel search jobs return, it aggregates the search result. It has two important contributions. First, it achieves super-linear speedup by reducing IO overhead. Second, it is the first parallel BLAST tool to provide exact e-value statistics in contrast to approximate e-value statistics of other contemporary parallel implementations of NCBI BLAST.

mpiBLAST’s exact e-value statistics requires two steps. It first collects the necessary statistical parameters for the entire database by performing a pseudo-run of the blast engine against the global database. Once it has the global parameters, it passes the global parameters (such as whole database length *n*, the total number of sequences *N*) to the parallel search jobs against segmented databases. mpiBLAST modifies some functionalities of NCBI BLAST (blast.c, blastdef.h, blastkar.c, and blastutl.c) so that global parameters can be fed externally and that information can be used to calculate exact e-values.

For accurate e-value correction, mpiBLAST requires prior knowledge of the entire database.

#### 2.3.2 NOBLAST

NOBLAST (Lagnel et al. 2009) provides new options for NCBI BLAST. It offers a way to correct e-values when split databases are used and the results need to be aggregated. E-value computation requires knowledge about the entire database size, the number of sequences in the whole database *N* and the total length of the database *n*. Using the values *N*, *n* and Karlin-Altschul statistical parameters which are independent of database size, the e-value can be computed using Karlin-Altschul statistics. NOBLAST first computes the length adjustment using the knowledge about the complete original database, then it computes effective search space using length adjustment, and finally, it computes the e-value using effective search space.

In principle, NOBLAST takes a similar approach to mpiBLAST, as both provide global statistical parameters to the search jobs against a segmented database so that that exact e-value can be computed. While mpiBLAST’s main contribution is a parallel implementation and e-value correction comes from the need of producing the same output as the sequential counterpart, NOBLAST’s main contribution is an e-value correction. Both tools require prior knowledge about the entire database. Both tools were developed before new e-value statistics were introduced, so they didn’t address e-value corrections for the BLAST programs that use new statistics.

#### 2.3.3 Comparison of mpiBLAST and NOBLAST to Incremental BLAST

While the former two BLAST tools can provide exact e-value statistics when Karlin-Altschul statistics are used and the knowledge about the entire database is available a priori, they can’t be used when the database keeps changing or two different search results against two different instances of similar databases needs to be aggregated. Comparison among these three tools are outlined in table 2.

**Table 2:** Comparison among three different BLAST tools that explicitly deal with e-value statistics. Incremental BLAST supports e-value correction across time and space without prior knowledge of the entire database. The other tools can correct e-values in limited scenarios.

| Feature | mpiBLAST | NOBLAST | Incremental BLAST |
| --- | --- | --- | --- |
| E-value correction for Karlin-Altschul Statistics | ✓ | ✓ | ✓ |
| E-value correction for Spouge Statistics | ✗ | ✗ | ✓ |
| Aggregate search results against pre-planned database segments | ✓ | ✓ | ✓ |
| Aggregate search results against arbitrary database instances | ✗ | ✗ | ✓ |
| Reuse existing search results | ✗ | ✗ | ✓ |

**Table 3:** Fidelity of Incremental BLAST in three consecutive time periods.

| Period | Database | Database Size |  | Evalue Match | Hit Match |
| --- | --- | --- | --- | --- | --- |
|  |  | Current | Incremental |  |  |
| 0 | nt | 80,740,533,243 | 80,740,533,243 | 100% | 100% |
|  | nr | 17,686,779,866 | 17,686,779,866 | 100% | 100% |
| 1 | nt | 113,749,495,340 | 33,008,962,097 | 100% | 100% |
|  | nr | 23,752,080,639 | 6,065,300,773 | 100% | 100% |
| 2 | nt | 152,471,828,601 | 38,722,333,261 | 100% | 100% |
|  | nr | 30,030,148,449 | 6,278,067,810 | 100% | 100% |

**Table 4:** Fidelity of Incremental BLAST(blastp) in two consecutive time periods.

| Period | Database Size |  | E-value Match | Hit Match |
| --- | --- | --- | --- | --- |
|  | Current | Incremental |  |  |
| 0 | 40,077,622,077 | 40,077,622,077 | 100% | 100% |
| 1 | 162,267,258 | 19,192,851,238 | 100% | 100% |

## 3 Methods

### 3.1 e-value correction in incremental setting

Correct e-value computation requires actual database length in both Spouge statistics and Karlin-Altschul statistics. While split-database parallel blast applications like mpiBLAST and NOBLAST has prior knowledge about actual database length, the incremental blast setting uses current database length. The former passes the actual database length to each of their parallel jobs, thus forcing the statistics module to compute correct e-values from the beginning. In the incremental setting, whenever new data arrives in the database, the search is refined in two steps. First, the search is run on new sequences to the database. Second, results are merged from the saved “current” and new search results. Merging requires re-evaluation of the e-values for all hits and their corresponding High Scoring Pairs (HSPs) using the total database-length.

#### 3.1.1 e-value correction for Karlin-Altschul statistics

Let, current database length is *n_c_*, length of the newly arrived part is *n_d_*. Number of sequences in current and the newly arrived part of the database are *N_c_* and *N_d_*. So, Actual length of the updated database, *n_t_* = *n_c_* + *n_d_*. Total number of sequences in updated database, *N_t_* = *N^c^* + *N^d^*.

Actual query length *m* does not change with the change in database. We have to recompute effective length *l* by solving the fixed point equation for new database length using equation 6.

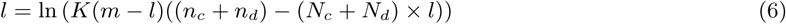

Once, we have the updated length adjustment *l*; we can either recompute e-values for all the matches or correct the e-values. To recompute all the e-values from scratch, we can use the formula 7.

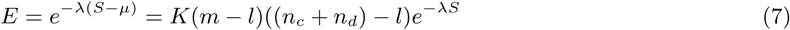

Alternatively, we can correct the e-values from current value. We will first use *l* to re-compute value of effective search space. We will use the newly computed effective search space to re-calibrate e-values for all the reported HSPs from current and delta search results. Let, part and total effective search spaces are *D_part_, D_total_*. Then, corrected e-value is *E_total_* = *E_part_* + *Ke^−λS^ ×* (*D_total_ − D_part_*).

Both approaches require a constant number of arithmetic operations, but the former approach requires fewer arithmetic operations.

#### 3.1.2 e-value correction for Spouge statistics

For Spouge statistics, value of area does not change since it is a function of query length, sequence length and Gumbel parameters. But, database scale factor does change, and we need to account for that. If actual database length for part and total databases are *n_part_* and *n_total_*, then

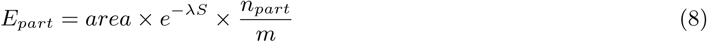

and

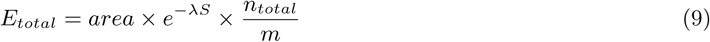

. So,

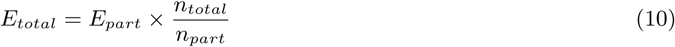

By these algebraic operations, we only have to re-scale the e-values instead of using Spouge’s e-value computation methods.

#### 3.1.3 e-value correction for extremely small values

Re-computing e-values for Karlin-Altschul statistics in Incremental BLAST always yields identical values as NCBI BLAST. In more than 99.9% cases, re-scaling e-values for Spouge statistics also yields identical values as NCBI BLAST. However, for some of the extremely small e-values reported by NCBI BLAST, Incremental BLAST reports 0.

We investigated the source of these mismatches. At time 0, NCBI BLAST rounded any e-values smaller than 1.0*e −* 180 to 0 which is also used by Incremental BLAST. Because the e-value for the same hit increases with database size, some of the e-values become greater than 1.0*e −* 180 and they can no longer be rounded to 0. NCBI BLAST computes e-values from the full database each time, so it is able to report correct changes in e-values with database size. In Incremental BLAST, re-scaling of the e-values would still report those e-values as 0 since it involves multiplying values between two time periods, and multiplying 0 with any number will still result in 0.

These e-values are straight forward to detect and we apply a fix by preventing NCBI BLAST programs from rounding small e-values to 0. We have modified the related portion of the code (*blast seqalign.cpp*) so that this approximation does not take place. With this approach, re-scaled e-values by Incremental BLAST should exactly match with the e-values reported by NCBI BLAST.

We comment out the first line and add the second line to implement this change in *blast seqalign.cpp*.

~~~
//double evalue = (hsp->evalue < SMALLEST_EVALUE) ? 0.0 : hsp->evalue; double evalue = hsp->evalue;
~~~

### 3.2 Merging two search results with correct e-value statistics

The hits reported by both searches are statistically significant. Once we correct e-values for both current search result and the new search result, we merge the hits into a single sorted list. Because Incremental BLAST reports some better scoring hits which NCBI BLAST misses, reporting only *max target seqs* hits will result in missing some of the lower scoring hits from NCBI BLAST. So, we store and report 2 *× max target seqs* hits. Algorithm 1 documents the procedure to merge the hits from two results for the same query. All the statistical parameters dependent on total databases size is recomputed to re-compute or re-scale the e-values. The hits are selected in the ascending order of their e-values (descending order of their scores).

#### Algorithm 1

Merging results for Karlin-Altschul/Spouge statistics

1: Input: *result*1*, result*2

*2: merged result ← Φ*

3: re-compute/re-scale e-values

4: *m, n ←* 0

5: **for** *i* = 1 *→ num of hits* **do**

6: *e − value*1*, score*1 *← min*(*result*1*.alignment*[*m*]*.hsps*)

7: *e − value*2*, score*2 *← min*(*result*2*.alignment*[*n*]*.hsps*)

8: **if** (*e − value*1 *< e − value*2) or *e − value*1 == *e − value*2 *and score*1 *> score*2) **then**

*9: merged result.add*(*result*1. *alignments[m]*)

10: *increment m*

11: **else**

12: *merged result.add*(*result*2*.alignments[n]*)

13: increment *n*

14: **end if**

15: **end for**

16: **return** *merged result*

More details on re-computing and re-scaling evalues is provided in A.1.

### 3.3 Incremental BLAST implementation

We develop Incremental BLAST as a lightweight software without modifying or re-implementing complex components of NCBI BLAST. We use most NCBI BLAST programs as black box routines. Users with NCBI BLAST+ installed in their system will simply need to install a Python script able to make use of existing NCBI BLAST+ programs. Any change in NCBI BLAST programs except e-value statistics will not require any change in Incremental BLAST software. Our tool will support multiple versions of NCBI BLAST if the BLAST+ programs use same statistics across these versions.

We show the software stack of Incremental BLAST in figure 2. Incremental BLAST software has three major components: User Interface, Incremental Logic, and Record Database. These modules interact with BLAST databases through BLAST+ programs.

**Figure 2:**
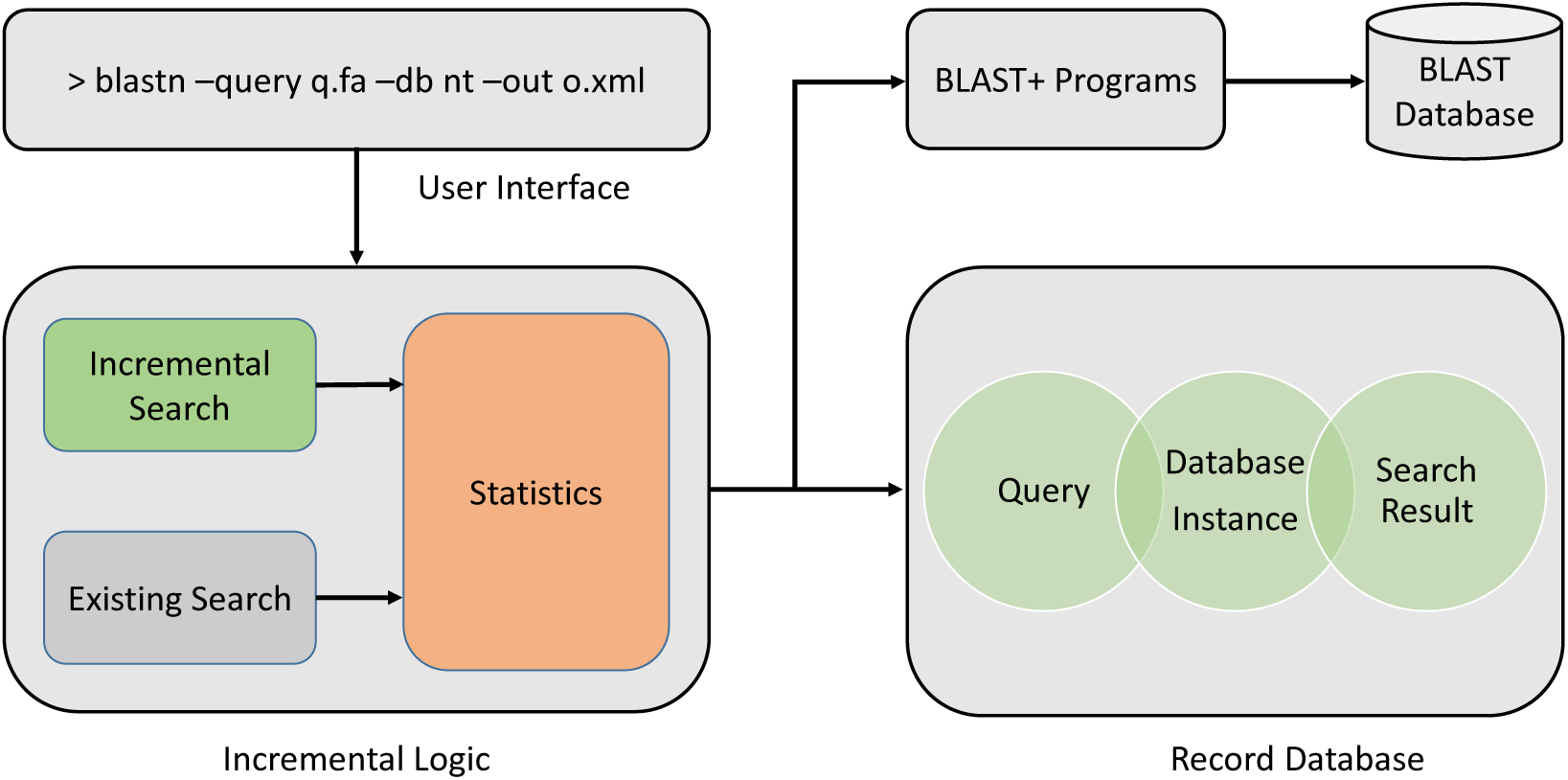
Software stack of Incremental BLAST. The user can initiate a search using the user interface. The search parameters are then passed to the Incremental Logic module. After performing an incremental search, this module corrects the e-values and merges the result. The incremental logic module looks into an external lightweight database module to decide whether and how to perform the incremental search. For actual search and incremental database creation, we use NCBI BLAST tools such as blastdbcmd, blastdbalias, blastp, blastn, etc.

**Command-line User Interface** In our current version, we provide a command line interface for Incremental BLAST which provides NCBI BLAST-like search options.

**Incremental Logic module** Incremental logic decides whether to perform a new BLAST search based on existing results. Whenever the user requests a new BLAST search, this module checks for any preexisting search result for this. If it does not find any preexisting result, it performs a regular NCBI BLAST. But, if there is a preexisting result, this module first compares the database instance from the time of past search with the present instance. If there is any difference in the size, this module builds a delta database by taking the sequences that differ in these two instances. Then it performs a new BLAST search only against the delta database and merges the old result with the new incremental result after statistical correction for e-values. Figure 3 shows different parts of this module. This allows multiple updates to current searches with little extra time investment. Incremental Logic module has three submodules: *Existing Search*, *Incremental Search*, and *Merge*.

**Figure 3:**
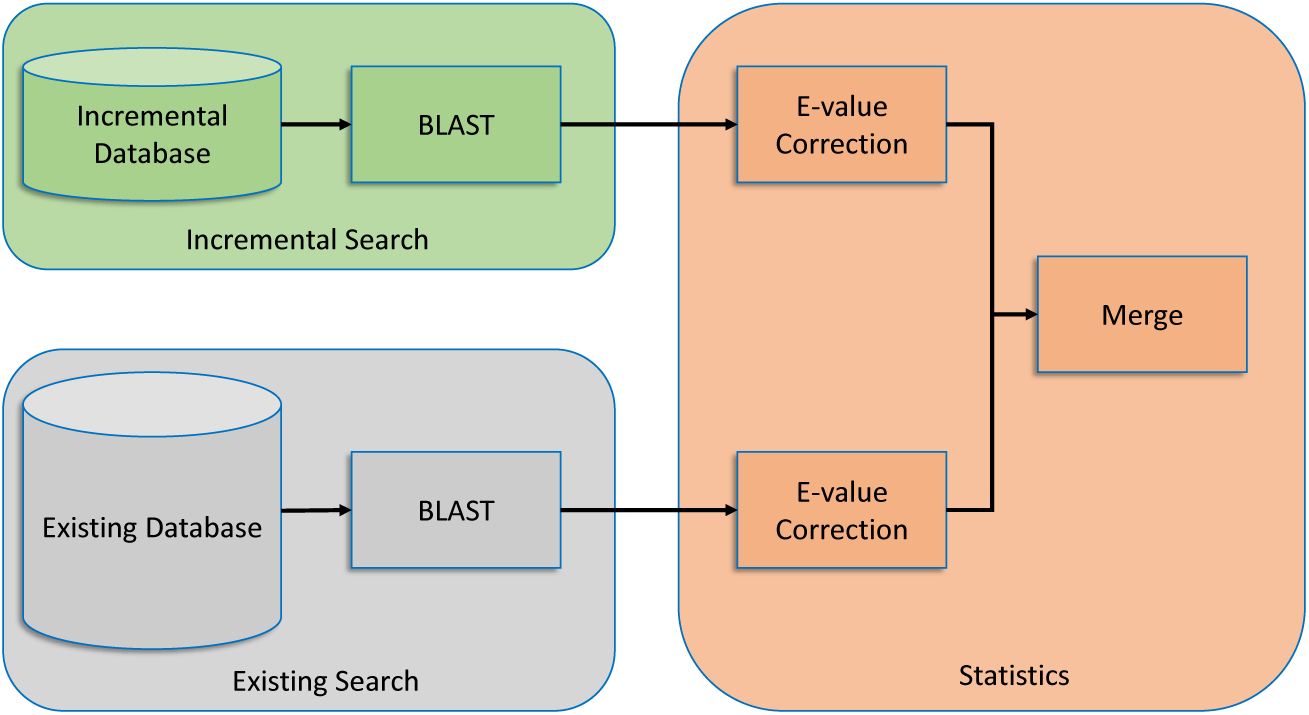
Sub-modules of Incremental Logic. Whenever the user initiates a BLAST job, Incremental Logic first checks if an existing search result is available. If there is a search result against an outdated BLAST database, a delta database consisting of newly added sequences are constructed. A BLAST search is then performed against the delta database(incremental database). In the final stage, the existing search result and the incremental search result are merged after correcting the e-values.

**Figure 4:**
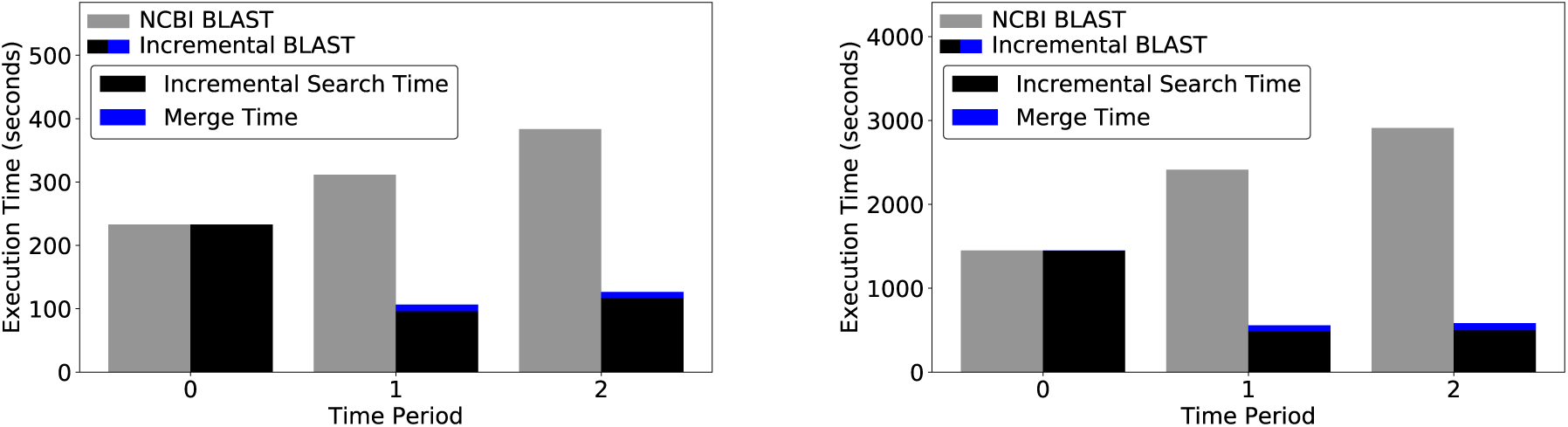
Performance for case study I

1. **Existing Search module** This sub-module looks up existing search result with the help of record database.
2. **Incremental Search Module** This sub-module constructs incremental database by comparing with past instance of the database and performs a BLAST search on the incremental database.
3. **Statistics module** This module reads the past search result and the new incremental search result, re-evaluates the e-values in both, and merge them according to their recomputed/re-scaled e-values.

The details of these three sub-modules are illustrated in figure 3.

**Record Database for storing incremental search results** Every time a BLAST search is performed, we save the instance of the database along with the search result in a lightweight SQLite database. We save a minimalistic index structure and size information that requires only a few bytes of storage. We keep the search parameters along with the search results as well.

### 3.4 Case studies

To demonstrate the efficiency and benefits of using the incremental blast program over standard NCBI BLAST, we analyze different actual nucleotide and protein sequence datasets as case studies.

#### 3.4.1 Case Study I: method verification

We explore the scenario where hits from a collection of 100 query sequences are updated to account for growth of NCBI sequence databases across the duration of the project. To demonstrate the the applications’s use for blast programs that use Karlin-Altschul statistics, we ran blastn against a nucleotide database (growing subsets of NCBI nt) for 100 nucleotide sequences from *Bombus impatiens* assembly available at ftp://ftp.ncbi.nlm.nih.gov/genomes/Bombus_impatiens/CHR_Un/bim_ref_BIMP_2.1_chrUn.fa.gz. To demonstrate its utility on blast programs that use Spouge statistics, we run blastp against a non-redundant protein database(growing subset of NCBI nr) for 100 protein sequences from *Bombus impatiens* assembly available at ftp://ftp.ncbi.nlm.nih.gov/genomes/Bombus_impatiens/protein/protein.fa.gz.

We demonstrate Incremental BLAST’s fidelity and performance over three time periods. The instances for nucleotide database changes through time as following:

1. **Time 0:** the nucleotide database comprises 52% of the full nt database. Both tool conduct search on the same database.
2. **Time 1:** the nucleotide database comprises 70% of nt. While NCBI BLAST searches 70% of nt, Incremental BLAST performs search only on 28% of nt. The database grew by 33% from time 0.
3. **Time 2:** the nucleotide database comprises 88% of nt. While NCBI BLAST searches 88% of nt, Incremental BLAST performs search only on 18% of nt. The database grew by 27% from time 1.

The instances for protein database changes through time as comprises 37%, 48%, and 60% of nt. As for nucleotide database search, Incremental BLAST performs search only on the incremental part and then it combine the new search result from earlier time period after e-value correction. The database grew by 32% and 25% in time 1 and 2 respectively from the earlier time periods.

More details on incremental database creation is provided in A.2. We compare the performance of NCBI BLAST and Incremental BLAST for each of these time periods.

#### 3.4.2 Case Study II: updating a query re-annotation of a novel transcriptomics dataset

Our second case study mimics a typical scenario in a transcriptome re-annotation project where a transcriptome is blasted after a certain period as a part of re-annotation pipeline. This uses a novel dataset not yet available on NCBI - a transcriptome of the venom gland of an oak gall wasp (see below) - and thus the identity of the assembled sequence was unknown and the sequence was not available to blast to itself. In casestudy I, we used existing query sequences from NCBI.

We conduct a blast search for the same query set for database instances in two time periods:

1. **Time 0:** the database comprises 70% of the non-redundant database nr (nr accessed on August, 2018). Both tools perform search on 70% of the database.
2. **Time 1:** the database comprises 100% of the non-redundant database nr. While NCBI BLAST performs search on 100% of nr, Incremental BLAST performs search on only 30% and reuses search result from time 0. Incremental BLAST merges the result from time 0 with the incremental search result after e-value correction.

We constructed these two database instances by combining database parts using blastdb aliastool packaged with BLAST+.

Given that the size of the transcriptome would take few months to complete on a single core, we ran this experiment with 640 cores distributed across 20 compute nodes, partitioning the 17927 queries into 20 query files and assigning each file per node. Given that each node will run a subset of queries against the same database, there is no need to re-compute the statistics for these results before we merge them. The time investment and results traditional and incremental approaches were compared.

#### 3.4.3 Case Study III: taxon-based incremental approach

Our third case study presents a special case of using a taxon-based incremental approach to obtain a fast, cost-effective and biologically relevant sequence similarity results. To achieve this goal, we examine the genes contained within an assembled transcriptome of the venom gland of a gall wasp of oak trees, the hedgehog gall wasp (***Acraspis erinacei***), a taxon lacking a closely related species with a genome. Gall wasps are a group of parasitic wasps that inject their eggs into plant tissues and through processes yet unknown, induce changes in plant development at the site of injection. These changes result in the construction of a niche for the gall wasp by inducing predictable modifications of plant tissues that both protect the wasp from the environment and feed the developing wasp. Genes important for inducing changes in the plant’s development are thought to be produced in the female venom gland during oviposition ((Vårdal 2006)). We performed separate blasts of the hedgehog gall wasp venom gland against transcriptomes of the closest relatives to gall wasps with curated genomes including three fairly equidistant taxa ((Peters et al. 2017)) - the parasitic wasp *Nasonia vitripennis*, the honey bee ***Apis mellifera***, and the ant ***Harpegnathos saltator***, - as well as the more distant model insect, ***Drosophila melanogaster***, upon which many insect gene annotations are based. We also blasted the transcripts to, an oak tree, ***Quercus suber***, to determine if some genes belonged to the host, and a model plant the soybean ***Glycine max***. A blastp search was conducted individually against each of the databases and results were merged using the statistics module of the Incremental BLAST. After this initial search, we then added to this analysis all remaining Hymenopteran species using Incremental BLAST, to assess the impact of adding more taxa on the top BLAST hits and further demonstrate the potential of incremental BLAST to add taxa progressively. We performed a blastp search result obtained from the merged database of those seven subsets to determine whether the same hits would have been found from our concatenated incremental analysis as from a combined single-instance run. These results were further compared with blastp results obtained by searching the complete nr database, thus allowing us to determine the extent of capture of the full dataset with this taxon subsampling approach.

#### 3.4.4 Data collection for case studies II and III

To obtain the venom gland transcriptome, 15 venom glands were dissected from newly emerged adult females from wild collected oak (oak spp.) hedgehog galls of *Acraspis erinacei* and placed in RNAlater. Pooled tissues were homogenized in lysis buffer using a Bead Ruptor 12 (Omni International) with additional lysis with a 26-gauge syringe. RNA was extracted from the sample using the RNaqueous Micro kit followed by DNase I treatment as specified by the kit and confirmed to be of good quality using the Bioanalyzer 2100 (Agilent). The Illumina HiSeq library was prepared from 200 ng RNA using the TruSeq Stranded mRNA kit and sequenced in 150 bp single-end reads across two Rapid Run lanes on the Illumina HiSeq 2500 (Penn State Genomics Core Facility, University Park) along with nine other barcoded wasp samples.

Raw sequence data (30.6 million reads) quality was assessed using FastQC v0.10.0 (Andrews & FastQC 2015), and appropriate trimming was conducted using Trimmomatic v0.35 (Bolger et al. 2014) with following parameters: ILLUMINACLIP:TruSeq3-PE.fa:2:30:10 LEADING:20 TRAILING:20 MINLEN:50 AVGQUAL:20. This procedure removed 0.12% of total reads. *de novo* transcriptome assembly from these QC-passed trimmed reads was performed on the Trinity RNA-Seq *de novo* Assembler (version: trinityrnaseq r20140717)((Grabherr et al. 2011)). The transcriptome assembly consists of 44, 440 transcripts with a contig N50 of 865 bases. Transdecoder v5.3.0 ((*TransDecoder (Find Coding Regions Within Transcripts)* 2018 (accessed September 15, 2018)) was used to predict 17927 likely protein sequences (adult venom gland trinity.fasta.transdecoder.pep) used for Case Study II and III.

## 4 Results

### 4.1 Incremental BLAST program

The Incremental BLAST program v1.0 includes a collection of python scripts which can be downloaded at https://bitbucket.org/sajal000/incremental-blast. The user needs to copy the source folder and run the following command from this directory to install Incremental BLAST:

~~~
./IncrementalBLAST-installer.sh
~~~

The python scripts are:

**Main program: iBLAST.py** This program provides incremental BLAST search options. It takes in a regular blast search command and performs incremental search. An example usage of the script is:

~~~
python iBLAST.py “blastp -db nr -query Trinity-tx.fasta -outfmt 5 -out result.xml”
~~~

**Merge Scripts** These scripts are used to merge two BLAST search results in xml format and produce an xml output with corrected e-values.

1. **BlastpMergerModule.py:** This is used to merge results obtained using Karlin-Altschul statistics (e.g., blastn results)
2. **BlastnMergerModule.py:** This is used to merge results obtained using Spouge statistics (e.g., blastp results)
3. **BlastpMegerModuleX.py** These scripts merge more than two blast results. They require number of results to merge, the input results and output.

~~~
python BlastpMergerModule.py input1.xml input2.xml output.xml
python BlastnMergerModule.py input1.xml input2.xml output.xml
python BlastpMergerModuleX.py 3 input1.xml input2.xml input3.xml output.xml
~~~

### 4.2 Case Study I: method verification

In case study I, we validate whether we can get the same results from a single NCBI BLAST as from the Incremental BLAST(3). In all three time periods, we find all the hits reported by NCBI BLAST for blastn were recovered by Incremental BLAST in the same order, including 3964 hits at time 0, 4150 hits at time 1, and 4924 hits at time 2, thus validates Incremental BLAST for Karlin-Altschul statistics. We observe similar results for blastp as well. In three time periods, Incremental BLAST reports the exact same hits in the exact same order as NCBI BLAST. The numbers of reported hits in these three time periods for blastp are 45154, 46, 356, and 46869. Reporting the same results for blastp validates Incremental BLAST for Spouge statistics.

At time period 1, out of 46, 356 e-values (taking one HSP from each hit with minimum e-value), only 21(0.045%) e-values were fixed using the method described in section 3.1.3. At time period 2, 0.132%(62 out of 46869 hits) e-values were fixed similarly.

For *δ* increase in database size Incremental BLAST performs (1 + *δ*)*/*(*δ*) times faster than NCBI BLAST. Figure 5 shows that for blastp and blastn Incremental BLAST achieves the speed-up as mentioned above in two different time periods.

**Figure 5:**
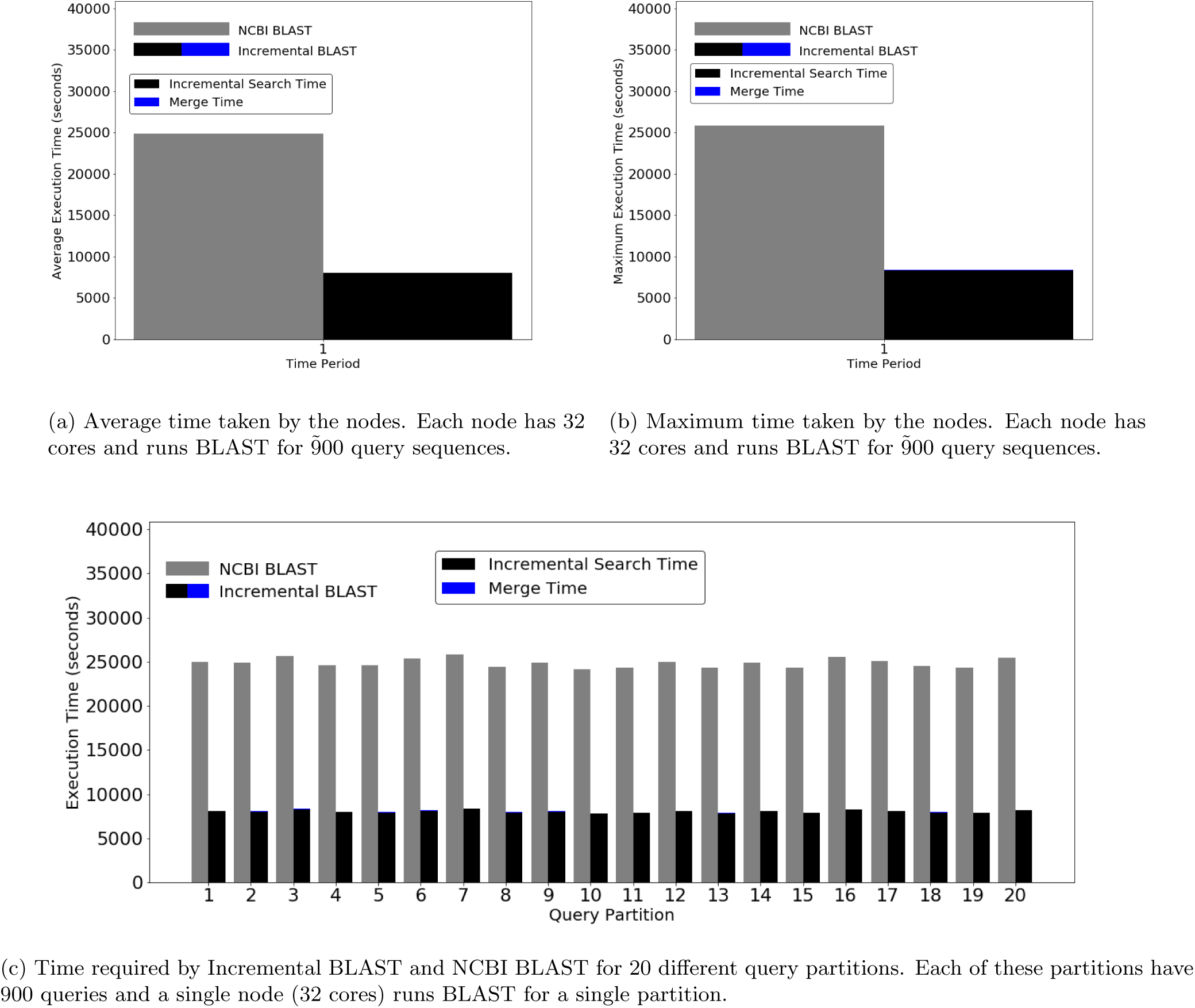
Performance for case study II. Time required by NCBI BLAST and Incremental BLAST. The average time taken is 24862 seconds(6 hours, 54 minutes) for NCBI BLAST, and 8009 seconds(2 hours, 14 minutes) for Incremental BLAST. Merge time for each of these task is 40 seconds on average. Maximum time for these three are 25835 seconds (7 hours, 11 minutes), 8334 seconds (2 hours, 19 minutes), and 49 seconds. By both account Incremental BLAST is 3.1 times faster than NCBI BLAST. This speedup complies with our projected speedup (1 + 0.48)*/*0.48 = 3.08.

### 4.3 Case Study II: Incremental BLAST is efficient for large alignment tasks on novel datasets

We perform search using Incremental BLAST and NCBI BLAST in two time periods with 48% increase in the nr database in between. In both time periods, Incremental BLAST reports the same hits with in the same order as NCBI BLAST, with 0.045% e-value mismatches in time period 1 after re-computing the e-values due to the issue discussed in casestudy I(4). We eliminate these mismatches by the approach described in section 3.1.3.

Figure 5 shows the time comparison between NCBI BLAST and Incremental BLAST. We see that for 48% increase in the database size, Incremental BLAST is 3.1% times faster than the NCBI BLAST in achieving the new result. The time needed for e-value correction and merging the results is minimal (less than a minute using only 20 cores).

### 4.4 Casestudy III: taxon-specific searches can expedite informatics

To examine the fidelity of Incremental BLAST when merging multiple (taxon-specific) databases (7), first, we compared the Incremental BLAST merged results from multiple individual BLAST(blastp) searches on seven biologically relevant taxa to results obtained when a BLAST search was performed against a database combining all the sequences belonging to these taxa. The result exhibits a high fidelity of 99.7% match (55 mismatches of 17927 when comparing the top hits for each query, 0.30679979918%). As in casestudy I and II, these mismatches are due to early approximation and can be eliminated by the method described in section 3.1.3. Then, we compared (presented in Table 5) the merged BLAST results of individual taxon-level database search with the BLAST results obtained in case-study II (time period 1) where the same queries were searched against the entire nr database to better understand relative time savings vs. accuracy of taxon-guided approaches. The taxon-specific approach is much more time-efficient and computationally inexpensive as it searched much smaller databases. With our initial set of 6 taxa we sampled only 0.35% of the nr database and retrieved 8.124% of the top hits obtained when searching nr. Although this number is low, the identity of top hits is likely to be similar even if the best taxonomic hit to that gene was not retrieved, as gall wasps do not have any close relatives sequenced but rather many equidistant relatives. Given this, we then added in the rest of Hymenoptera to see if this improves the number of shared top hits. With this analysis we BLAST-ed only 1.17% of the total nr yet obtained 87.75% similarity in top hits to a full nr BLAST. This demonstrates the potential of performing more taxon-guided approaches to save on the costs of large-scale BLASTs. Performing the analysis this way also enabled improved curation of hits by taxon.

**Table 5:** Potential for taxon-guided searches. Comparison of merged BLAST results from multiple individual blast search with a separate BLAST search conducted against a completed nr database, showing that biologically relevant taxa can be added incrementally to obtain similar results to nr with much fewer queries.

| NCBI<br>Taxon ID | Species | % nr sequences<br>covered | Number of nr<br>top hits covered | % nr top hits<br>covered |
| --- | --- | --- | --- | --- |
| 7425 | *Nasonia vitripennis* (jewel wasp) | 0.02% | 853 | 4.84% |
| 7460 | *Apis mellifera* (honey bee) | 0.02% | 207 | 1.17% |
| 10380 | *Harpegnathos saltator* (Jerdon's jumping ant) | 0.03% | 347 | 1.96% |
| 7227 | *Drosophila melanogaster* (fruit fly) | 0.08% | 6 | 0.034% |
| 58331 | *Quercus suber* (cork oak) | 0.09% | 0 | 0.00% |
| 3847 | *Glycine max* (soybean) | 0.11% | 22 | 0.12% |
| 7399 | Rest of Hymenoptera | 0.83% | 14281 | 80.98% |
| **Total** | **multiple** | **1.17%** | **15476** | **87.75%** |

### 4.5 Incremental BLAST finds better scoring hits missed by NCBI BLAST

While Incremental BLAST finds all the hits reported by NCBI BLAST in the same relative order, Incremental BLAST reports some better scoring hits that NCBI BLAST misses for all the case studies. Since case study II covers the most number of hits, we quantified these missed hits for this case study. NCBI BLAST misses 1.57% (13171 out of 837942 top hits) of the better scoring hits. Command line NCBI BLAST uses a search parameter *max target seqs* in an unintended way where instead of reporting all the *best max target seqs* hits, it has a bias toward *first max target seqs* hits Shah et al. (2018). In this process, it misses some of the better scoring hits that is discovered in a later phase of the search A.3. This is an extra advantage of Incremental BLAST over NCBI BLAST. Since the former works on smaller databases and then combines the results instead of searching a single large database, it has more candidate hits to choose from for reporting final hits.

## 5 Discussion

In this paper, we have introduced Incremental BLAST, an incremental local alignment tool that can provide identical results as NCBI BLAST. Our tool is capable of performing the same task as NCBI BLAST at a much improved speed by adding to previous search results, an appraoch enabled by the development of novel statistical methods of e-value correction. This aproach can be used to combine multiple search results performed at different times or with different groups (e.g. taxa), thus facilitates novel ways of performing sequence alignment tasks and incorporating domain knowledge. For *δ* fraction increase in the database size, Incremental BLAST can perform (1 + *δ*)*/δ* times faster than NCBI BLAST (i.e. 10% growth in database size will yield 11 times speedup for Incremental BLAST). It should be noted that for small increase in the database size (which is most likely scenario between two searches), Incremental BLAST provides large speedup factor.

While Incremental BLAST provides almost identical results as NCBI BLAST, there are a small number of mismatches caused by early approximations of very low e-value values (high scores) which we fix through small modification in NCBI BLAST code. There is another source of mismatch that proves Incremental BLAST more successful in discovering better hits. While Incremental BLAST finds 100% of the hits that NCBI BLAST reports in the same order, it reports many high scoring hits that NCBI BLAST misses due to an early approximation used by NCBI BLAST’s heuristic search algorithm.

With the expansion of genetic data available in NCBI, computational time is becoming ever more burdensome, resulting in analyses that take months to complete with substantial financial cost. This problem is aggravated by cheaper sequencing technology leading to ever-larger transcriptomics projects with substantially more samples to analyze. Our program can help relieve the cost burden. It enables iterative updates for re-annotation of genome and transcriptome assemblies, useful given rapid changes in the nr databases across the duration of a project. Specific datasets of interest can be added to previous searches, such as new genome releases or large phylogenetic studies. As demonstrated in Case Study III, the program can be applied to transcriptomic or metagenomics projects by merging the results of knowledge-guided blasts only on groups that are biologically relevant. This enables iterative exploration by taxon and facilitates curation of blast results.

## Acknowledgements

We like to thank Andy Deans and István Mikó for their contribution in data collection. This work was supported by *ICTAS II* @ Virginia Tech. Computations for this research were performed on Virginia Tech’s *Advanced Research Computing (VT ARC)* and Pennsylvania State University’s *Institute for CyberScience Advanced CyberInfrastructure (ICS-ACI)*.

## A Supplementary Materials

### A.1 e-value correction

We use algorithm 2 to re-compute e-values for BLAST programs using Karlin-Altschul statistics. We first aggregate database sizes from two input results and use the aggregated size*N* to compute length adjustment *l*. Using *N* and *l*, we re-compute e-values for both results.

#### Algorithm 2

Recomputing e-values for Karlin-Altschul Statistics

1: Input: *result*1*, result*2

2: *n ← result*1*.n* + *result*2*.n*

3: *m ← result*1*.m*

4: *N ← result*1*.N* + *result*2*.N*

5: *l ← recompute length adjustment*(*n, m, N*)

6: *recompute evalues*(*result*1*, l, N*)

7: *recompute evalues*(*result*2*, l, N*)

We use algorithm **??** to re-scale e-values for BLAST programs using Spouge statistics. First we aggregate the database sizes for two input results, and scale the e-values by a factor of the ratio between aggregate database size and the individual database size.

#### Algorithm 3

Rescale e-values for Spouge Statistics

1: *Input: result*1, *result2*

2: *db length*1 ← *result*1.*db length*

3: *db length*2 ← *result*2.*db length*

4: *db length ← db length*1 + *db length*2

5: *rescale evalues*(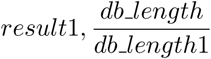)

6: *rescale evalues*(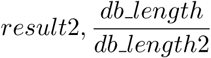)

### A.2 Creating experimental databases

Pre-formatted BLAST databases such *nt* and *nr* come in incremental parts. With progression of time, new sequences are packaged in parts and added to the databases.

#### A.2.1 Databases for casestudy I

For case study I, we consider three time steps when the nt and nr databases had 30, 40, and 50 parts. For these three time periods, we construct three databases as instances of nt and nr by combining 30, 40, and 50 parts using BLAST tool *blastdb aliastool*. The incremental databases between two periods are also constructed. Table 6 shows different instances of *nt* databases in three different periods.

**Table 6:** Incremental nt Databases for case study I.

| Period | Database parts | Number of sequences | Number of bases | Longest sequence length |
| --- | --- | --- | --- | --- |
| 0 | 0-29 | 25,117,275 | 80,740,533,243 | 774,434,471 bases |
| 0-1 | 30-39 | 8,389,596 | 33,008,962,097 | 275,920,749 bases |
| 1 | 0-39 | 33,506,871 | 113,749,495,340 | 774,434,471 bases |
| 1-2 | 40-59 | 8,891,258 | 38,722,333,261 | 129,927,919 bases |
| 2 | 0-49 | 42,398,129 | 152,471,828,601 | 774,434,471 bases |

Table 7 shows different instances of *nr* databases in three different periods.

**Table 7:** Incremental nr databases for casestudy I

| Period | Database parts | Number of sequences | Number of residues | Longest sequence length |
| --- | --- | --- | --- | --- |
| 0 | 0-29 | 49,468,463 | 17,686,779,866 | 36,507 residues |
| 0-1 | 30-39 | 15,878,318 | 6,065,300,773 | 35,523 residues |
| 1 | 0-39 | 65,346,781 | 23,752,080,639 | 36,507 residues |
| 1-2 | 40-49 | 16,448,075 | 6,278,067,810 | 38,105 residues |
| 2 | 0-49 | 81,794,856 | 30,030,148,449 | 38,105 residues |

**Table 8:** Incremental nr databases for casestudy II

| Period | Database | Number of sequences | Total residues | Longest sequence length |
| --- | --- | --- | --- | --- |
| 0 | 0-63 | 109,407,071 | 40,077,622,077 | 38,105 residues |
| 0-1 | 64-90 | 52,860,187 | 19,192,851,238 | 74,488 residues |
| 2 | 0-90 | 162,267,258 sequences | 59,270,473,315 | 74,488 residues |

#### A.2.2 Databases for casestudy II

We construct nr database instances for time 0 and 1 by combining 64 and 90 parts respectively. We combine these parts using *blastdb aliastool*.

### A.3 Explanation for NCBI BLAST missing many top hits

Due to the early cutoff of max target sequence used by its heuristic algorithm. NCBI BLAST performs search in two phases. In earlier phase(ungapped extension), it starts with matching a seed substring between target and query sequence and then extends the matching pair in both direction without allowing any gap. In this phase, BLAST algorithm assigns some scores to these matching pairs and keeps only the very high scoring pairs using a cutoff determined by e-value cutoff or number of maximum hits. In the gapped phase, these selected high scoring pairs are further extended in both directions while allowing gaps and these evolved pairs get changed socres. Some of the pairs that did not make the cut during the ungapped extension, can become high scoring pairs. For a larger database, these missed opportunities are higher in number because there are more potential pairs in the ungapped phase. Since Incremental BLAST is combining results from smaller databases, it misses relatively smaller number of those high scoring hits compared to NCBI BLAST.

